# Dedifferentiation and neuronal repression define Familial Alzheimer’s Disease

**DOI:** 10.1101/531202

**Authors:** Andrew B. Caldwell, Qing Liu, Gary P. Schroth, Douglas R. Galasko, Shauna H. Yuan, Steven L. Wagner, Shankar Subramaniam

## Abstract

Early-Onset Familial Alzheimer’s Disease (EOFAD) is a dominantly inherited neurodegenerative disorder elicited by over 300 mutations in the *PSEN1, PSEN2*, and *APP* genes^1^. Hallmark pathological changes and symptoms observed, namely the accumulation of misfolded Amyloid-β (Aβ) in plaques and Tau aggregates in neurofibrillary tangles associated with memory loss and cognitive decline, are understood to be temporally accelerated manifestations of the more common sporadic Late-Onset Alzheimer’s Disease. The complete penetrance of EOFAD-causing mutations has allowed for experimental models which have proven integral to the overall understanding of AD^2^. However, the failure of pathology-targeting therapeutic development suggests that the formation of plaques and tangles may be symptomatic and not describe the etiology of the disease^3,4^. In this work, we used an integrative, multi-omics approach and systems-level analysis in hiPSC-derived neurons to generate a mechanistic disease model for EOFAD. Using patient-specific cells from donors harboring mutations in *PSEN1* differentiated into neurons, we characterized the disease-related gene expression and chromatin accessibility changes by RNA-Seq, ATAC-Seq, and histone methylation ChIP-Seq. Here, we show that the defining disease-causing mechanism of EOFAD is dedifferentiation, causing neurons to traverse the lineage-defining chromatin landscape along an alternative axis to a mixed-lineage cell state with gene signature profiles indicative of less-defined ectoderm as well as non-ectoderm lineages via REST-mediated repression of neuronal lineage specification gene programs and the activation of non-specific germ layer precursor gene programs concomitant with modifications in chromatin accessibility. Further, a reanalysis of existing transcriptomic data from *PSEN1* patient brain samples demonstrates that the mechanisms identified in our experimental system recapitulate EOFAD in the human brain. Our results comprise a disease model which describes the mechanisms culminating in dedifferentiation that contribute to neurodegeneration.

## Results

Non-Demented Control (NDC), *PSEN1*^*M146L*^, and *PSEN1*^*A246E*^ hiPSCs were generated and differentiated into neurons as previously described^5,6^ (Fig. 1A, Fig. S1A). Subsequent RNA-Seq and differential expression analysis of magnetically-purified neurons identified 10020 differentially expressed genes (DEGs) in *PSEN1*^*M146L*^ neurons and 4093 DEGs in *PSEN1*^*A246E*^ neurons relative to NDC with substantial overlap between the two mutations (Fig. 1B, Fig. S1B-C). In order to characterize the functional programs modulated and to delineate the transcriptional regulators controlling them, we first carried out Panther^7^ enrichment analysis using the Reactome Pathway and Gene Ontology (GO) databases, revealing consistent upregulated enrichment of gene sets related to cell cycle, pluripotency, inflammation, and non-ectoderm lineage dedifferentiation (EMT, ECM organization, and mesendoderm lineage specification captured by cardiac system-related development gene sets) combined with consistent downregulation of gene sets related to neuron lineage specification and neuron function in both *PSEN1*^*M146L*^ and *PSEN1*^*A246E*^ neurons (Fig. 1C-D). Pre-ranked Gene Set Enrichment Analysis (GSEA)^8^ using the Hallmark database confirmed positive enrichment of cell cycle and inflammatory programs (Fig. S1D). As terminally differentiated neurons are generally thought to exist in the quiescent G_0_ phase, yet *PSEN1*^*M146L*^ and *PSEN1*^*A246E*^ neurons exhibit significant enrichment for cell cycle-related processes, we sought to determine the neuron cell cycle state. This revealed that *PSEN1*^*M146L*^ and *PSEN1*^*A246E*^ neuron populations have a significantly lower percentage of cells in the G_0_ and S phases and a higher percentage in the G2/M phases, suggesting they have re-entered the cell cycle (Fig. 1E). To validate these results, we performed qPCR in *PSEN1*^*M146L*^, *PSEN1*^*A246E*^, and *PSEN1*^*H163R*^ hiPSC-derived neurons for key cell cycle related and REST-repressed (*GAD1*) genes, recapitulating expression patterns observed in the *PSEN1*^*M146L*^ and *PSEN1*^*A246E*^ RNA-Seq data (Fig. S1E). As miRNAs have previously been shown to regulate neuronal lineage specification, cell cycle, and dedifferentiation processes^9,10^, to characterize the transcriptional regulation of these enriched gene sets we first used the miRTarBase database to perform pre-ranked GSEA and Enrichr^11^ miRNA target enrichment in the *PSEN1*^*M146L*^ and *PSEN1*^*A246E*^ conditions, demonstrating that the targets of functionally-relevant miR-124 and miR-9 are substantially upregulated in both conditions (Fig. 1F, Fig. S1F). To determine whether this is due to loss of miR-124 and miR-9, we performed miRNA-qPCR, revealing that miR-124 and miR-9 levels are significantly decreased in *PSEN1*^*M146L*^ and *PSEN1*^*A246E*^ neurons relative to NDC (Fig. 1G). To further characterize the transcriptional regulation of these altered gene programs, we performed two complementary enrichment approaches: TF-target enrichment by Enrichr and pre-ranked GSEA and motif analysis by ISMARA^12^ to characterize the TFs and miRNAs whose motif activity most significantly changes between the *PSEN1*^*M146L*^ or *PSEN1*^*A246E*^ and NDC conditions. Enrichr analysis using the ENCODE/ChEA consensus database revealed enrichment for TFs related to cell cycle (E2F family members, FOXM1)^13^ and pluripotency (SOX2, NANOG, KLF4)^14,15^ amongst the upregulated genes and TFs related to neuronal specification (REST, CREB1, NRF1)^16–18^ amongst the downregulated genes (Fig. 1H). Pre-ranked GSEA using the ENCODE/ChEA database as well as a custom TF database of AD disease ontology-associated TFs showed analogous positive enrichment of cell cycle and pluripotency regulators identified by Enrichr as well as TFs related to inflammation (IRF1, RelA, IRF8)^19,20^, dedifferentiation (YAP1/TAZ, SOX9, SNAI2, NOTCH1, SALL4)^21–23^, and neuronal specification (RFX2, MYT1L, NeuroD1)^24,25^ (Fig. 1I). Furthermore, the top transcriptional regulators identified by TF-target and ISMARA motif enrichment analysis coalesced into the same six general categories of disease-causal endotypes identified by ontology enrichment analysis; four upregulated (pluripotency, cell cycle, inflammation, and dedifferentiation) and two downregulated (neuronal specification, lineage miRNA) (Fig. 1J). The ISMARA score for the majority of transcriptional regulators across all categories and both mutations, which describes the predicted gain or loss of activation, followed the direction of log_2_ fold change of regulator gene expression. Interestingly, miR-124 and miR-9 were identified by ISMARA in the *PSEN1*^*M146L*^ and *PSEN1*^*A246E*^ neurons as having decreased activity and are known repressors of the most significantly activated TF identified, REST^26^.

**Fig. 1.**
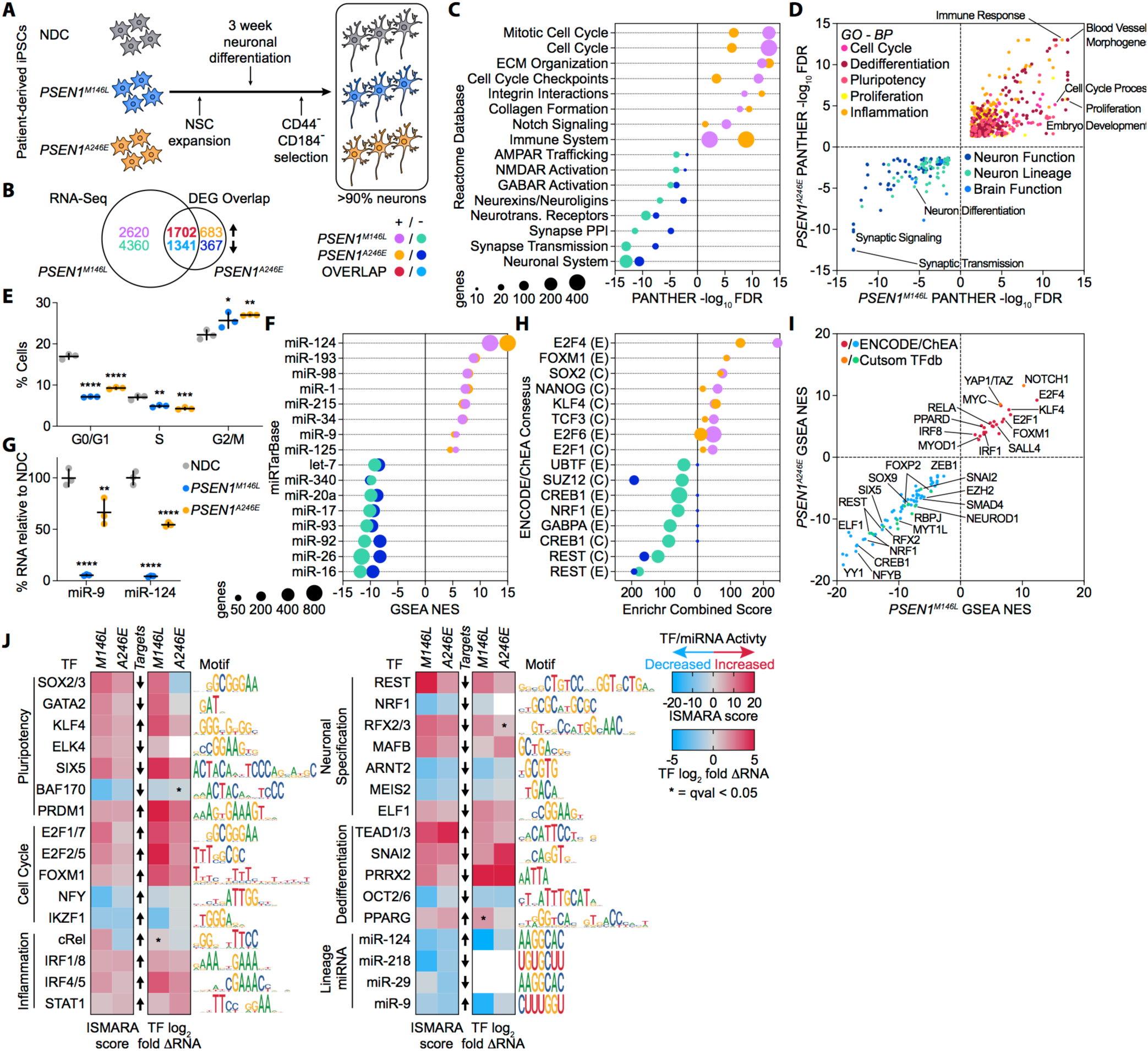
PSEN1 hiPSC-derived neurons undergo dedifferentiation through the activation and repression of key endotypes. **A** Patient-derived Non-Demented Control (NDC), *PSEN1*^*M146L*^, and *PSEN1*^*A246E*^ hiPSCs were differentiated into neurons and purified by CD44^−^/CD184^−^ selection. **B** Overlap of differentially expressed genes (DEGs) in *PSEN1*^*M146L*^ and *PSEN1*^*A246E*^ hiPSC-derived neurons relative to non-demented control (NDC) with a False Discovery Rate (FDR) adjusted p-value (q-value) < 0.01 and log_2_ fold-change > [0.5]. (*n* = 4) **C** PANTHER functional enrichment of DEGs in *PSEN1*^*M146L*^ and *PSEN1*^*A246E*^ hiPSC-derived neurons using the Reactome Pathway database. (FDR p-value < 0.05) **D** Common PANTHER functional enrichment scores of DEGs in *PSEN1*^*M146L*^ and *PSEN1*^*A246E*^ hiPSC-derived neurons using the Gene Ontology – Biological Process database, curated into ontologically related, disease-associated cellular programs (endotypes). (FDR p-value < 0.05) **E** Percentage of NDC, *PSEN1*^*A246E*^, or *PSEN1*^*M146L*^ hiPSC-derived neurons in each cell cycle phase; cells stained with FxCycle™ Far Red Stain and assayed by flow cytometry; p-value: * = < 0.05, ** = < 0.01, *** = < 0.001, **** = < 0.0001; mean ± SD. (*n* = 3) **F** Top positively and negatively enriched miRNA by Normalized Enrichment Score (NES) in the *PSEN1*^*M146L*^ and *PSEN1*^*A246E*^ conditions by pre-ranked Gene Set Enrichment Analysis (GSEA) using the miRTarBase v7.0 miRNA database (FWER < 0.01). **G** Total miRNA levels of miR-9 and miR-124 in *PSEN1*^*M146L*^ and *PSEN1*^*A246E*^ hiPSC-derived neurons relative to NDC. p-value: ** = < 0.01, **** = < 0.0001; mean ± SD. (*n* = 3) **H** Enrichr enrichment of upregulated (purple, *PSEN1*^*M146L*^; orange, *PSEN1*^*A246E*^) and downregulated (green, *PSEN1*^*M146L*^; dark blue, *PSEN1*^*A246E*^) genes using the ENCODE/ChEA Consensus TF database (see methods). **I** TFs with commonly enriched targets by pre-ranked GSEA in the *PSEN1*^*M146L*^ and *PSEN1*^*A246E*^ conditions using the ENCODE/ChEA Consesus TF or Custom neural TF database. **J** ISMARA motif analysis in *PSEN1*^*M146L*^ and *PSEN1*^*A246E*^ hiPSC-derived neurons identifying the TFs and miRNAs with most significant motif-associated activity change, based on ISMARA z-score and Pearson correlation, curated into 6 key modulated disease-associated endotypes.

Next, we aimed to evaluate whether these observed transcriptional changes were due to modulation of the chromatin state. To this end, we performed ATAC-Seq on NDC and *PSEN1*^*M146L*^ hiPSC-derived neurons, identifying 14223 regions of chromatin with differential accessibility between the two conditions (Fig. 2A-B), 5640 of which occurred within DEGs (Fig. 2C, Fig. S2). Metascape^27^ ontological analysis of the differentially accessible promoter (pATAC) and putative enhancer (eATAC) regions uncovered significant enrichment of GO-BP and Reactome gene sets related to cell cycle, non-ectoderm lineage and dedifferentiation, inflammation, and neuron specification across all four types of chromatin state change (increase or decrease in accessibility of promoter or enhancer regions) (Fig. S3A). In order to identify the TF motifs enriched in these regions of differential chromatin accessibility, we first performed Homer motif analysis^28^, finding significant enrichment of pluripotency (SOX2, OCT4) and lineage definition (BORIS, SOX9)^29,30^ motifs within regions of increased accessibility and enrichment of neuronal (NRF1, ELF1, ELK1)^31^ motifs within regions of decreased accessibility (Fig. 2D). HINT TF footprinting analysis^32,33^ identified TFs with similar ontological function, predicting increased TF activity of precursor and lineage regulators (OCT1/3/4, SOX10, TWIST1)^34^ and decreased TF activity of neuron regulators (NRF1, CREB1, FOXP2, FOXO6)^35–37^ and EMT repressors (OVOL2)^38^ (Fig. 2E). Strikingly, ATAC read density and motif accessibility by HINT around NRF1 sites is substantially decreased in the *PSEN1*^*M146L*^ condition relative to NDC (Fig. 2F). Enrichr TF-target enrichment of genes with differentially accessible chromatin and a significant change in expression identified many of the same transcriptional regulators found by ATAC-Seq motif analysis as well as *PSEN1*^*M146L*^ and *PSEN1*^*A246E*^ neuron RNA-Seq TF-target and ISMARA motif analysis, including regulators for cell cycle (E2F4, FOXM1), pluripotency (SOX2, OCT4, NANOG) and dedifferentiation (SALL4) among the upregulated genes and TFs related to neuron lineage specification (REST, PRC2 components)^39^ and neuron mitochondrial function (NRF1, CREB1) among the downregulated genes (Fig. 2G-H). These same endotype-associated changes are observed by ontological enrichment analysis with Metascape, demonstrating that the increased gene expression and enrichment observed for cell cycle, dedifferentiation, and pluripotency programs and decreased gene expression and enrichment observed for neuron lineage and function is influenced by the chromatin accessibility state (Fig. 2I-J). Metascape and Enrichr analysis of DEGs grouped by the location of differential chromatin accessibility revealed key insights at the functional and transcriptional regulatory level; interestingly, genes with a decrease in differential promoter accessibility were significantly enriched for cell cycle programs, but the median expression of this subset of genes was increased in the *PSEN1*^*M146L*^ condition (Fig. S3B). Whereas enrichment of neuron lineage specification genes is observed for genes with differential promoter and putative enhancer accessibility, later stage neuronal gene programs, such as axonogenesis, synaptic signaling, and behavior are primarily enriched within the decreased enhancer accessibility group (Fig. S3B-C). Furthermore, while NRF1 and CREB1 target genes were enriched among the downregulated genes with differential chromatin accessibility occurring within the promoter, REST and PRC2 target genes were enriched among the downregulated genes with differential chromatin accessibility occurring outside the promoter (Fig. S3D-E).

**Fig. 2.**
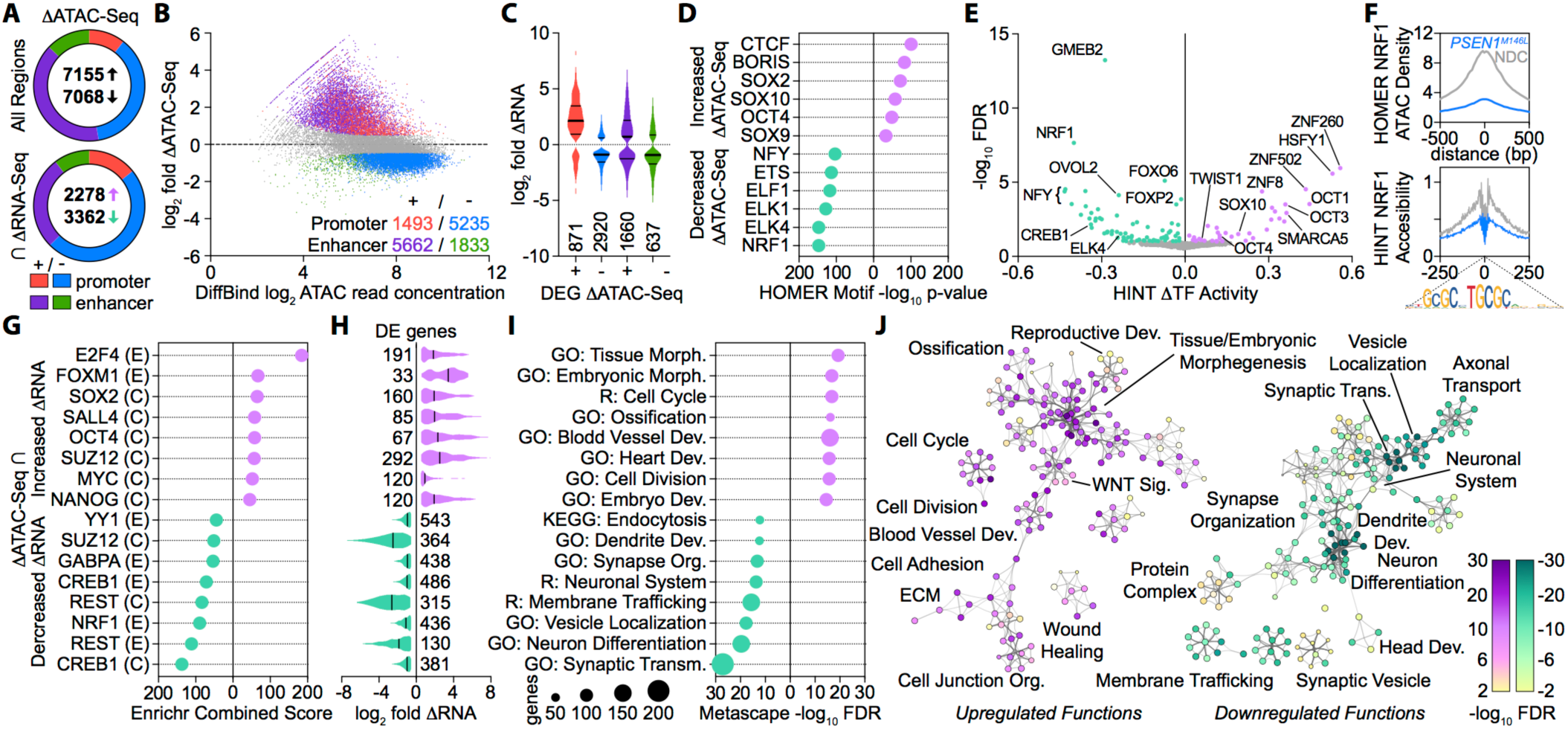
Modulation of chromatin accessibility drives differential gene expression and dedifferentiation in PSEN1^M146L^ hiPSC-derived neurons. **A** Differentially accessible regions of chromatin in *PSEN1*^*M146L*^ hiPSC-derived neurons relative to NDC as measured by ATAC-Seq; top, all differentially accessible regions; bottom, differentially expressed genes with differentially accessible regions of chromatin. Colors maintained A-C. (*n* = 2) **B** MA Plot of differential chromatin accessibility occurring within the promotor region or outside the promotor region. **C** Violin plot of RNA log_2_ fold-change for genes with differential accessibility and gene expression. **D** Homer TF motif enrichment of all regions with increased accessibility (top, purple) or decreased accessibility (green, bottom). **E** TF motifs with a significant (adj. p-value < 0.1) change in activity relative to NDC by HINT TF footprinting analysis. **F** Homer ATAC-Seq read density (top) and HINT TF accessibility (bottom) in *PSEN1*^*M146L*^ and NDC hiPSC-derived neurons around NRF1 motifs. **G** Enrichr enrichment using the ENCODE/ChEA Consensus TF database of genes with differential accessibility and increased gene expression (top, purple) or decreased gene expression (bottom, green). **H** Violin plot of RNA log_2_ fold-change for targets of enriched TFs in G. **I-J** Metascape enrichment of genes with ATAC-Seq and increased gene expression (purple) or decreased gene expression (green) **I** ranked by FDR p-value and **J** visualized as enrichment networks.

As chromatin accessibility can be modulated through changes in histone methylation^40^, and TF enrichment revealed REST and the histone-modulating PRC2 complex target genes as being significantly downregulated, we performed ChIP-Seq on NDC and *PSEN1*^*M146L*^ hiPSC-derived neurons for the activating mark H3K4Me3 and the repressive mark H3K27Me3. This revealed 7309 regions of differential H3K4 trimethylation and 21230 regions of differential H3K27 trimethylation (Fig. 3A). To investigate the potential transcriptional regulators associated with these methylation changes, we performed Homer motif analysis, identifying motifs for TFs related to cell cycle (E2F1, E2F4, E2F7), pluripotency (OCT4-SOX2-Nanog, SP1) and neuron lineage (REST) (Fig. 3B-C). Further, Metascape ontology analysis revealed non-ectoderm lineage gene sets enriched amongst genes with increased H3K4Me3 or decreased H3K27Me3 as well as neuron lineage and synaptic function gene sets enriched amongst genes with decreased H3K4Me3 or increased H3K27Me3 (Fig. 3D-E). Of these differentially methylated regions, 1583 upregulated genes intersected with increased H3K4Me3 and 667 with decreased H3K27Me3, whereas 733 downregulated genes intersected with decreased H3K4Me3 and 1147 intersected with increased H3K27Me3 (Fig. 3F-G, Fig. S4A-B). Enrichr TF enrichment of these differentially expressed genes with differential methylation revealed that the upregulated genes are enriched for control by pluripotency-related TFs (SOX2, NANOG, KLF4), the dedifferentiation TF SALL4, and the PRC2-complex factor SUZ12, while the downregulated genes are enriched primarily for REST and PRC2-complex factors SUZ12 and EZH2 (Fig. 3H). Metascape analysis of these subset of genes revealed similar upregulation of non-ectoderm lineage development and downregulation of synaptic function processes in genes with differential H3K4Me3 or H3K27Me3 (Fig. 3I). Next, we integrated the ATAC-Seq data with the histone methylation ChIP-Seq data, revealing that the intersection of differentially expressed genes with differential methylation concomitant with differential chromatin accessibility (Fig. 3J) showed a substantial overlap. Moreover, the genes with increased H3K4Me3 or decreased H3K27Me3 as well as increased chromatin accessibility and expression are enriched for gene sets related to EMT and dedifferentiation gene sets and transcriptional control by lineage factors such as GATA2, OCT4, and SOX2, while the genes with decreased H3K4Me3 or increased H3K27Me3 as well as decreased chromatin accessibility and expression are enriched for gene sets related to neuron lineage and synaptic function and transcriptional control by REST and PRC2 complex factors (Fig. 3K, Supplementary Fig.4C). Read profiles of previously identified AD-associated genes (*VGF*)^41^ as well as non-ectoderm lineage (*TEAD2, TCF7L2*)^42^ or neuroectoderm lineage (*MYT1L*) specifying genes show chromatin accessibility and histone methylation changes which correspond directionally to their increased (Fig. 3L) or decreased (Fig. 3M) gene expression. Unsurprisingly, the locus around the two key neuron lineage miRNAs downregulated in both *PSEN1*^*M146L*^ and *PSEN1*^*A246E*^ conditions, miR-9 and miR-124, show decreased chromatin accessibility and H3K4Me3 as well as increased H3K27Me3 (Fig. 3N). A heatmap corresponding to directionally correlated RNA expression, promoter accessibility, and H3K trimethylation for an expanded list of genes belonging to the core gene programs identified here or previously identified as AD marker genes illustrates key gene expression changes occurring due modifications of the chromatin landscape (Fig. 3O). In order to further delineate the mechanistic gene changes occurring due to modifications at the chromatin level, we separated the genes into categories of those with a change in chromatin accessibility concomitant with differential H3K4 or H3K27 trimethylation or those with a change in promoter chromatin accessibility alone; Enrichr TF and Metascape ontology enrichment revealed that genes in the former group are primarily involved in pluripotency, dedifferentiation, and neuron lineage specification gene programs, while those in the latter group are involved in cell cycle, chromatin conformational modification, mitochondrial, and vesicle transport gene programs (Fig. S4D-G). As REST is transcriptionally upregulated and predicted to be activated as evidenced by ISMARA analysis and target gene repression, we carried out REST ChIP-Seq in *PSEN1*^*M146L*^ neurons. Here, we identified 1379 REST-bound target genes, a substantial portion of which are shared with other previously-reported REST target lists (Fig. S5A-B)^43,44^. Furthermore, ontology enrichment analysis of these REST genes revealed significant enrichment for the canonical REST processes neuron differentiation and synapse development, as well as dedifferentiation endotype-associated processes, such as blood vessel development (Fig. S5C). Overall, this multi-omics, integrative analysis illustrates that REST and PRC2 neuronal target genes are transcriptionally repressed due to a change in chromatin accessibility driven by modification of H3K27Me3 and H3K4Me3 status, whereas NRF1 and CREB1 neuronal target genes are repressed due to loss of chromatin accessibility in the promoter without change in the H3K4Me3 or H3K27Me3 methylation state.

**Fig. 3.**
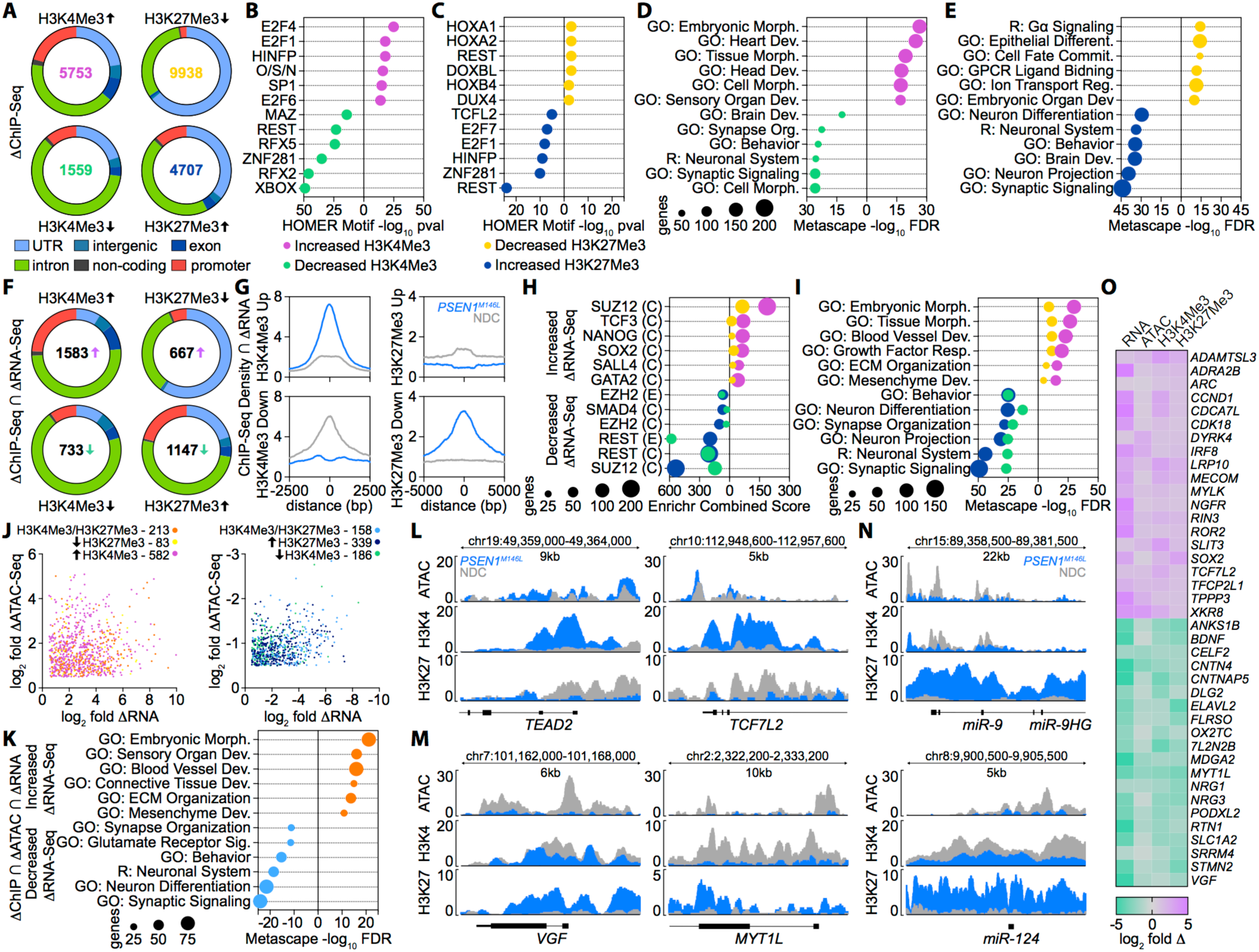
Dedifferentiation in PSEN1^M146L^ hiPSC-derived neurons is caused by changes in histone methylation leading to modulation of chromatin accessibility. **A** Differentially methylated histone regions in *PSEN1*^*M146L*^ hiPSC-derived neurons relative to NDC as measured by ChIP-Seq for H3K4Me3 or H3K27Me3. (*n* = 1) **B-C** Homer TF motif enrichment of all regions with a significant increase or decrease in **B** H3K4Me3 or **C** H3K27Me3 (O/S/N = OCT4-SOX2-Nanog). **D-E** Metascape functional enrichment of all regions with a significant increase or decrease in **D** H3K4Me3 or **E** H3K27Me3. **F** DEGs with a directionally corresponding increase or decrease in H3K4Me3 or H3K27Me3. **G** Enrichr enrichment of DEGs with significant change in H3K4Me3 and H3K27Me3 using the ENCODE/ChEA Consensus TF database. **H** ChIP-Seq read profiles around the summits of differential H3K4Me3 or H3K27Me3 methylation for DEGs with significant change in H3K4Me3 or H3K27Me3. **I** Metascape enrichment of DEGs with significant change in H3K4Me3 or H3K27Me3. **J** log_2_ fold change in chromatin accessibility and RNA expression in genes with (left) increased gene expression and corresponding change in H3K trimethylation state or (right) decreased gene expression and corresponding change in H3K trimethylation state. **K** Metascape enrichment of genes with differential H3K trimethylation, chromatin accessibility, and gene expression. **L-N** ATAC-Seq, H3K4Me3, and H3K27Me3 read profiles around the promoter regions of **L** genes with increased expression, **M** genes with decreased expression, or **N** miRNAs with decreased expression. **O** Heatmap of log_2_ fold change in RNA expression (RNA), chromatin accessibility within the promoter (ATAC), H3K4Me3 status, or H3K27Me3 (projected in negative log_2_ space to correspond to expression) status for endotype-associated or previously identified AD-associated genes.

Patient-specific hiPSC-derived neurons are a powerful model system to investigate EOFAD-causing mutations in a mechanistic, systems-level fashion; in order to validate whether the mechanisms identified in *PSEN1*^*M146L*^ neurons faithfully capture the disease-related processes occurring in AD patients, we reanalyzed a previous transcriptomic study^45^ (GSE39420) of posterior cingulate cortex brain tissue from NDC and *PSEN1*^*M139T*^ EOFAD donors (Fig. 4A). Expression analysis revealed a substantial number of DEGs in *PSEN1*^*M139T*^ brain overlapping with *PSEN1*^*M146L*^ and *PSEN1*^*A246E*^ neurons, particularly in the case of the downregulated genes (Fig. 4B). Interestingly, many uniquely DEGs in *PSEN1*^*M139T*^ patient brain shared similar ontology with those from *PSEN1*^*M146L*^ and *PSEN1*^*A246E*^ neurons (Fig. S6A-B). This shared ontology in *PSEN1*^*M139T*^ patient brain was similarly observed by Metascape (Fig. SC-D) and pre-ranked GSEA (Fig. S6E) analysis for all six key disease-associated endotypes identified. Furthermore, the DEGs uniquely found in *PSEN1*^*M139T*^ patient brain are enriched for the same ontological gene sets as *PSEN1*^*M146L*^ and *PSEN1*^*A246E*^ neurons, demonstrating a shared mechanism and functional change between *PSEN1*^*M139T*^ brain and *PSEN1*^*M146L*^ and *PSEN1*^*A246E*^ neurons independent of common DEGs (Fig. S6F). Enrichr TF/miRNA-target enrichment and ISMARA motif analysis showed similar TFs and miRNAs related to neuronal differentiation (REST, PRC2 complex, miR-124), non-ectoderm lineage specification and dedifferentiation (PRRX2, SMAD4, miR-124), pluripotency (SOX2, GATA2, LEF1), and cell cycle (E2F1/7, miR-9) as observed in *PSEN1*^*M146L*^ and *PSEN1*^*A246E*^ neurons, with REST identified as the top enriched/activated TF in *PSEN1*^*M139T*^ brain by both approaches (Fig. 4C-D, Fig. S6G). Further, comparison of ISMARA and pre-ranked GSEA results for TF motifs and TF targets, respectively, for the six core endotype gene programs identified here revealed consistent TF modulation between *PSEN1* mutant neurons and *PSEN1*^*M139T*^ brain (Fig. 4E). The activation of transcriptional regulators mechanistically driving these endotype changes was also identified by Ingenuity Pathway Analysis^46^, with directionally similar activation z-scores commonly predicted as upstream regulators for many key endotype-associated TFs across the three conditions (Fig. S6H). This consistent enrichment similarity between *PSEN1* mutant neurons and *PSEN1*^*M139T*^ brain is also seen at the gene set level by Panther enrichment and pre-ranked GSEA approaches, particularly for the downregulated synaptic signaling and plasticity gene sets (Fig. 4F). As the transcriptional regulator MYT1L was identified at the intersection of our multi-omics approach as a key downregulated gene and recently was shown to promote neuronal differentiation and repress non-ectoderm lineage^24^, we performed pre-ranked GSEA on a consensus list of MYT1L-regulated genes^24,47^. Strikingly, the normalized enrichment scores for MYT1L were similar across all three disease conditions, suggesting that loss of MYT1L, a REST-repressed and miR-124-promoted gene, contributes to loss of the neuronal state (Fig. S6I).

**Fig. 4.**
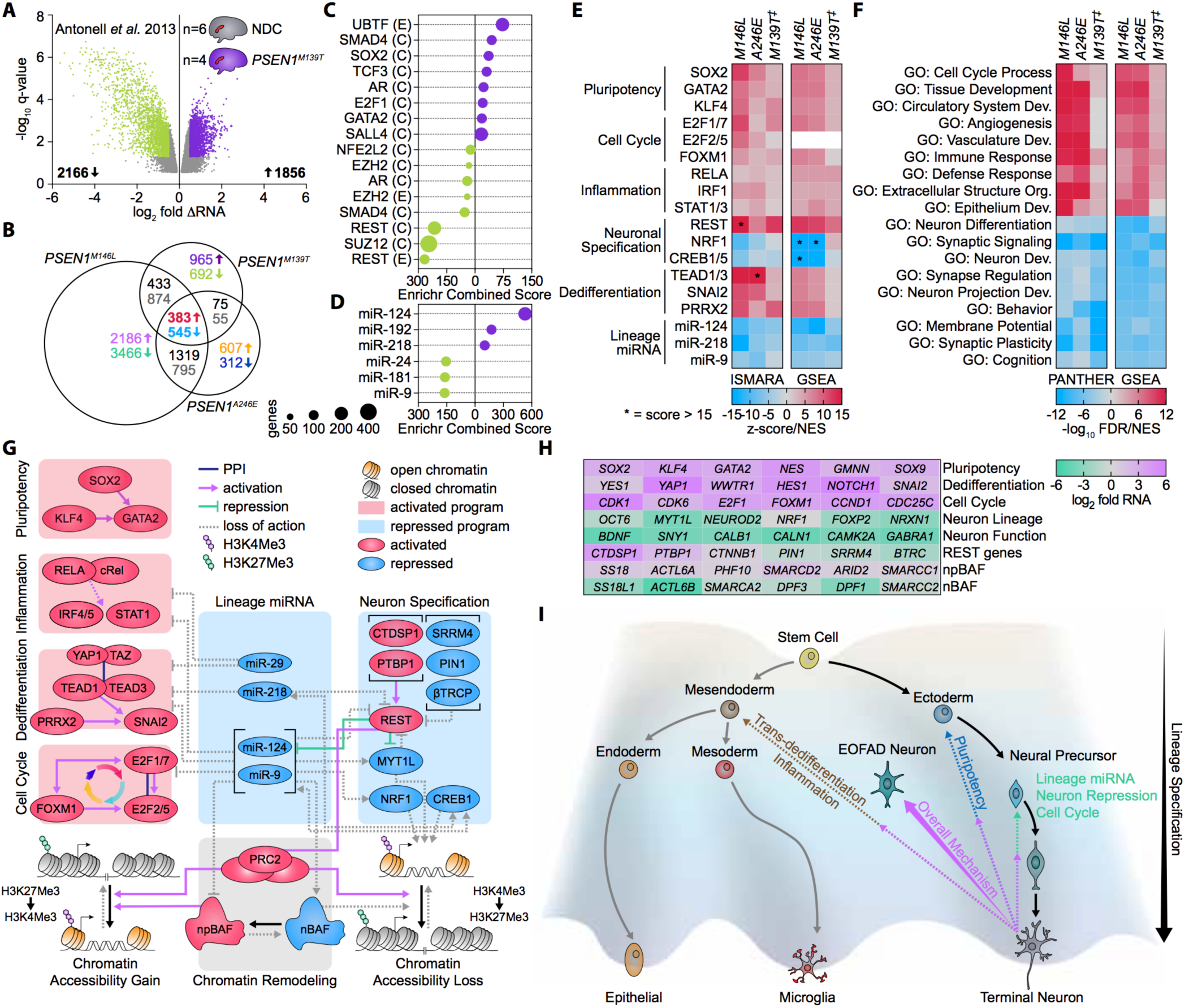
Mechanisms of neuronal dedifferentiation occurring in PSEN1 hiPSC-derived neurons are observed in EOFAD patient brains. **A** log_2_ fold change of differentially expressed genes in *PSEN1*^*M139T*^ human brain samples relative to non-demented control (NDC) with a False Discovery Rate (FDR) adjusted p-value (q-value) < 0.05. (NDC *n* = 6, *PSEN1*^*M139T*^ *n* = 4) **B** Venn-Euler diagram of DEGs in common between *PSEN1*^*M139T*^ human brain samples and *PSEN1*^*M146L*^ or *PSEN1*^*A246E*^ hiPSC-derived neurons. **C-D** Enrichr enrichment of upregulated and downregulated DEGs in *PSEN1*^*M139T*^ human brain samples using the **C** ENCODE/ChEA Consensus TF database or **D** miRTarBase v7.0 miRNA database. **E** Heatmap of ISMARA motif analysis (left) or pre-ranked GSEA TF enrichment (right) directional scores indicating gain or loss of activity for transcriptional regulators related to six key endotypes in *PSEN1*^*M139T*^ human brain (‡) samples and *PSEN1*^*M146L*^ or *PSEN1*^*A246E*^ hiPSC-derived neurons. **F** Top differentially enriched Gene Ontology – Biological Process gene sets by Panther (left) or pre-ranked GSEA (right) enrichment in *PSEN1*^*M139T*^ human brain samples and *PSEN1*^*M146L*^ or *PSEN1*^*A246E*^ hiPSC-derived neurons. **G** Mechanistic integration of the six disease-associated endotypes and key transcriptional regulators leading to remodeling of the chromatin state and dedifferentiation in the EOFAD disease state. **H** Heatmap of endotype-associated DEGs in the *PSEN1*^*M146L*^ condition. **I** A lineage specification landscape model. Repression of neuron specification and neuron lineage miRNAs concomitant with activation of cell cycle, pluripotency, inflammation, and dedifferentiation programs revert EOFAD neurons along an alternative axis into a less differentiated, precursor state with non-ectoderm lineage characteristics.

## Discussion

In this work, we used a patient-specific hiPSC-derived neuron model system and an integrative multi-omics approach to characterize the mechanistic changes occurring at the transcription factor and chromatin dynamics level leading to transcriptional dysregulation in EOFAD due to *PSEN1* mutations. Expanded enrichment analysis of *PSEN1* mutant neuron transcriptomics, combined with chromatin dynamics analysis in *PSEN1*^*M146L*^ neurons, revealed six modulated gene programs integral to the understanding of the overall disease mechanism: pluripotency, dedifferentiation, cell cycle re-entry, inflammation, lineage miRNA, and neuronal specification encompassing lineage definition and synaptic function. Strikingly, the key aberrant gene programs in *PSEN1*^*M146L*^ and *PSEN1*^*A246E*^ neurons driving neurodegeneration are analogously dysregulated in *PSEN1*^*M139T*^ human patient brains. The comparative enrichment scores for the top transcriptional regulators belonging to these programs across the EOFAD mutations gives insight into the documented severity of each, as *PSEN1*^*M146L*^ causes a particularly early onset and accelerated progression of the disease.

The integration of ATAC-Seq and histone methylation ChIP-Seq with TF target and binding site enrichment analysis of RNA-Seq data in *PSEN1*^*M146L*^ neurons revealed that transcriptional modulation of the core gene programs have distinct mechanisms underlying the chromatin accessibility changes; whereas upregulation of pluripotency genes and REST-mediated repression of neuronal genes occur due to promoter-proximal changes in H3K4Me3 and H3K27Me3 states leading to changes in accessibility, the downregulation of NRF1 and CREB1-mediated neuronal maturation and mitochondrial function genes primarily occur due chromatin accessibility changes within promoter regions potentially driven by alternative histone modifications combined with the loss of the pioneering TF capabilities of NRF1 and CREB1^48^.

However, these core gene programs and the transcriptional regulators orchestrating their modulation do not exist in isolation from one another, and in certain key aspects are intertwined in how they reshape the chromatin landscape (Fig. 4G-H). For example, commitment to a neuron lineage involves the coordinated repression of REST, repression of the REST-recruited, histone-modifying complex PRC2, and switching of the chromatin-modulating SWI-SNF complex components from a npBAF (neuronal precursor) to a nBAF (neuronal) state, which predominantly occurs through increased expression of the CREB1-activated miRNAs miR-124 and miR-9^9,26,49^. Further, miR-124 and MYT1L have been shown to restrict commitment to an neuroectoderm lineage through their repression of non-ectoderm lineage genes, while miR-9 plays an integral role in the cell cycle exit required for neuronal differentiation through its repression of key cell cycle regulators upon activation^10^. These multifaceted functions of REST, miR-124, miR-9, and MYT1L highlight their overarching identity as key neuron lineage determination factors; therefore, the combined reactivation of REST and downregulation of miR-124, miR-9, and MYT1L cause EOFAD neurons to dedifferentiate along an alternative axis through the repression of neuronal differentiation concomitant with activation of trans-dedifferentiation programs (Fig. 4I). Consequently, the resulting EOFAD neurons exist in a precursor-like state with transcriptional signatures of ectoderm and non-ectoderm lineage alike.

While early-onset AD accounts for at most 5% of all cases, it undoubtedly shares the hallmark pathological signatures and symptoms with the late-onset, sporadic form of the disease. However, the similarities between these two forms of AD with respect to the mechanisms driving neurodegeneration occurring before the onset of pathology and cognitive decline are less well known. A previous analysis of over 600 sporadic AD and non-demented, age-controlled subject brains identified EMT, a non-ectoderm dedifferentiation process significantly enriched in our *PSEN1*^*M146L*^ and *PSEN1*^*A246E*^ neurons as well as the *PSEN1*^*M139T*^ brains, as the top key upregulated gene program which facilities the progression of healthy aging into neurodegenerative AD^50^. Interestingly, miR-124 is the top enriched transcriptional regulator for the disease-contributing gene programs identified in this previous study, and we found substantial gene overlap between the EMT genes dysregulated in sporadic AD patients and *PSEN1* mutant neurons presented here (Fig. S6J). Moreover, the NIH AMP-AD program’s Agora list of sporadic AD susceptibility genes identified by multiple omics approaches show an overlap with EOFAD at both the gene and gene program level, with key drivers and markers of neuronal dedifferentiation, including *REST, HDAC1, ELAVL4, GABRA4, SCN2A*, and *VGF* similarly identified (Fig. S6K). This commonality of cellular programs altered in both the early-onset familial and late-onset sporadic forms offers a key insight into the underlying mechanistic basis for neurodegeneration in AD and provides a basis for novel therapeutic intervention at an earlier state in the progression of the disease.

## Acknowledgments

The authors thank Dr. Kristen Jepsen and the UC San Diego Institute for Genomics Medicine for their assistance with the sequencing performed in this research.

## Author contributions

A.B.C. performed the ChIP-Seq and ATAC-Seq experiments, carried out the data processing and systems analysis of the RNA-Seq, ChIP-Seq, and ATAC-Seq experiments, developed the mechanistic framework, and wrote the manuscript with contributions to the Methods section from other authors; A.B.C. and S.S. were involved in the characterization of the mechanisms and networks associated with EOFAD; S.H.Y. was involved in the design of hiPSC differentiation into neurons and the validation experiments along with S.L.W. and S.S.; Q.L. supervised by S.H.Y. carried out the hiPSC-neuron preparation, RNA isolation, qPCR validation, and cell cycle assay experiments; G.P.S. contributed to the RNA-, ChIP-, and ATAC-Seq; D.R.G. was involved in part of the study design; S.S. was involved in the overall study design, analysis, and revisions of the manuscript.

## Data and materials availability

All sequencing presented in this work is available at the NCBI GEO under the SuperSeries accession GSE123998, encompassing RNA-Seq (GSE123891), ATAC-Seq (GSE123997), and ChIP-Seq (GSE123884) datasets.

## Materials and Methods

### hiPSC line generation

Non-Demented Control (NDC), *PSEN1*^*M146L*^, and *PSEN1*^*H163R*^ fibroblasts were derived from skin biopsies in accordance with UC San Diego IRB approval, whereas *PSEN1*^*A246E*^ fibroblasts were obtained commercially (Coriell Cat. AG06840). The generation and characterization of the hiPSC lines were carried out as previously reported^5,51^ from fibroblasts by retroviral transduction using the reprogramming factors *OCT4, SOX2, KLF4*, and *C-MYC*. These four iPSC lines were used for all downstream neuron differentiation applications.

### Human neuron preparation

The protocol used for neuron preparation was previously described^6^. Briefly, neural stem cells (NSCs) were expanded in NSC growth medium (DMEM /F12, Glutamax™ (Thermo Fisher Cat. 10565018), 1x B-27 (Thermo Fisher Cat. 17504044), 1x N-2 (Thermo Fisher Cat. 17502001), 1x Penicillin-Streptomycin (Thermo Fisher Cat. 15070063), and 20ng/mL human bFGF-2 (BioPioneer Cat. HRP-0011)). At 80% confluence, the medium was changed to neural differentiation medium (DMEM /F12, Glutamax™, 1x B-27, 1x N-2, and 1x Penicillin-Streptomycin) for 3 weeks of differentiation. After differentiation, the cultures were dissociated with Accutase (Sigma Cat. A6964). Cells were resuspended in 200μL of IMag buffer (1x neural differentiation medium, 0.5µM EDTA, 0.5% Bovine Serum Albumin) followed by incubation with PE Mouse Anti-Human CD184 and CD44 antibodies (BD Biosciences Cat. 561733 and 561858, respectively) for 15min on ice in the dark. The mixture was washed with IMag buffer and subsequently incubated with magnetic beads (BD Biosciences) for 30min at room temperature. Magnetic bead separation was carried out for 8min according to the manufacturer’s protocol (BD Biosciences). The supernatant containing purified CD184^−^/CD44^−^ neurons was then removed and spun down for downstream applications.

### RNA Sequencing and Data Processing

For RNA-Seq, total RNA from magnetically purified human NDC, *PSEN1*^*M146L*^, and *PSEN1*^*A246E*^ (replicates, *n* = 4) hiPSC-derived neurons was prepared using RNeasy Plus Micro Kit (Qiagen Cat. 74034) according to the manufacturer’s protocol. On-column DNAse digestion was subsequently performed on total RNA extracts according to the manufacturer’s recommendations to remove any genomic contamination (Qiagen Cat. 79254). Libraries were prepared for RNA-Seq using the TruSeq Stranded Total RNA Library prep kit (Illumina, Cat. RS-122-2303) by the Ribo-Zero ribosomal RNA reduction method (Illumina, Cat. MRZG12324). Samples were sequenced at the UC San Diego Institute for Genomics Medicine (IGM) sequencing core on an Illumina HiSeq4000 generating Paired-End, 100bp reads with an average of 25 million reads per sample (Illumina, Cat. FC-410-1001).

RNA-Seq data preprocessing was performed using the BBDuk package (v37.95) in the BBTools suite^52^, removing sequencing adaptors and selecting for all paired-end reads above a quality score threshold (Phred Q>10). Trimmed RNA-Seq reads were mapped to the GRCh38.p12 human transcriptome using Salmon v0.9.12^53^ with the options --gcBias --seqBias -- biasSpeedSamp 5, followed by transcript level summation to the gene level using the R package tximport v1.8.0^54^. Differential expression analysis was performed for *PSEN1*^*M146L*^ and *PSEN1*^*A246E*^ iPSC-derived neurons relative to NDC using the R package DESeq2 v1.20.0^55^, selecting for genes with a q-value (FDR-adjusted p-value) of <0.01 and a log_2_ fold-change of >0.5. These lists of differentially expressed genes (DEGs) for the *PSEN1*^*M146L*^ and *PSEN1*^*A246E*^ conditions were used for all subsequent analysis.

### ATAC-Sequencing and Data Processing

ATAC-Seq transposition experiments were performed as previously reported^56^ on 50,000 NDC and *PSEN1*^*M146L*^ (replicates, *n* = 2) hiPSC-derived neurons, using the Illumina Nextera DNA Sample Preperation kit (Illumina, Cat. 15028523) and Qiagen MinElute PCR Purification kit (Qiagen, Cat. 28004). ATAC-Seq libraries were generated from transposed DNA using the Kapa Biosystems Real-Time Library Amplification kit (Kapa Biosystems, Cat. 07959028001) according to the manufacturer’s recommendations, monitoring amplification by qPCR and stopping the reaction when all samples reached a fluorescence amplification intensity between standards 1 and 3. ATAC-Seq libraries were further purified using the Qiagen MinElute PCR Purification kit and sequenced at the UC San Diego IGM sequencing core on an Illumina HiSeq4000 platform generating Paired-End, 50bp reads with an average of 25 million reads per sample.

ATAC-Seq data preprocessing was performed using the BBDuk package (v37.95) in the BBTools suite, removing sequencing adaptors and selecting for all paired-end reads above a quality score threshold (Phred Q>10). Trimmed reads were aligned to the GRCh38 human genome (GCA_000001405.15 with no alternative analysis) using BBMap v37.95 in the BBtools suite with the options maxindel=20 ambig=random, followed by sorting and indexing of bam files using Samtools v1.3^57^, and annotation of PCR duplicates using the Picard v2.3.0 markDuplicates function with the option VALIDATION_STRINGENCY=LENIENT. All duplicates and mitochondrial, Chromosome X, Chromosome Y, and EBV reads were removed using Samtools v1.3 command view with the options -b -h -f 3 -F 4 -F 8 -F 256 -F 1024 -F 2048. To determine regions of open chromatin, i.e. those accessible by the Tn5 transposase, bam files were first converted to bedGraph format using bedtools v2.25.0^58^ functions bamtobed and genomecov, and then finally converted to bigWig format using UCSC bedGraphToBigWig. HMMRATAC v1.0^59^ was used to call peaks on the ATAC-Seq data and determine regions of open chromatin with the options –m 50,200,400,600 --score all. These specifically called open regions of chromatin were then passed to the R package *Diffbind* v2.8.0^60^ to determine regions of differential accessibility between the NDC and *PSEN1*^*M146L*^ conditions using the edgeR method. Differentially accessible regions of DNA were annotated using the R packages *ChIPseeker*^61^ and *TxDb.Hsapiens.UCSC.hg38.knownGene* v3.4.0^62^, defining the promoter region -1000 to 1000 bp from the TSS.

In order to determine the transcription factor footprints associated with the gain or loss of chromatin accessibility, we used HINT v0.11.1^32,33^ for the following steps; rgt-hint function footprinting with the options --atac-seq --paired-end --organism=hg38 to identify TF footprints in each sample; rgt-motifanalysis function matching with the options --organism=hg38 --motif-dbs ∼/hocomoco to match footprints to known TFs in the HOCOMOCO v11^63^ human pwm database; and rgt-hint function differential with the options --organism=hg38 --output-profiles and --window-size 200. Differential HINT footprinting was performed for all differentially accessible regions with corresponding differential gene expression over the entire HMMRATAC-called open chromatin region. In order to generate histograms of ATAC-Seq read density and H3K4Me3 or H3K27Me3 ChIP-Seq read density at the promoter-located summits of differential chromatin accessibility intersecting with differential gene expression, the Homer v4.10 function annotatePeaks was used with the options -size 2000 -hist 20 for ATAC-Seq read density, -size 5000 -hist 50 for H3K4Me3 read density, or -size 10000 -hist 100 for H3K27Me3 read density.

### ChIP-Sequencing and Data Processing

For H3K4Me3 and H3K27Me3 ChIP-Seq experiments, NDC and *PSEN1*^*M146L*^ (*n* = 1) hiPSC-derived neurons were processed using the Active Motif ChIP-IT High Sensitivity kit according to the manufacturer’s recommendations (Active Motif, Cat. 53040). Briefly, cells were washed with cold 1x PBS and collected in a 1.5 mL tube in PBS, followed by formaldehyde fixation for 10 min at room temperature. After addition of stop solution, cells were incubated at room temperature for 5 min and subsequently washed with 1x PBS Wash Buffer and resuspended in Chromatin Prep Buffer. Following incubation on ice for 10 min, cells were lysed by performing 30 plunges in a chilled dounce homogenizer. Samples were then resuspended in Active Motif ChIP Buffer and chromatin sonication performed using a probe tip sonicator on ice for a total of 10 min per sample (30x 20 sec intervals). Optimal sonicated chromatin fragment lengths were confirmed by agarose gel and used for IP reactions. 500 ng of purified, sonicated chromatin was incubated with 5 µL of H3K4Me3 (Active Motif, Cat. 39159), H3K27Me3 (Active Motif, Cat. 39155), or 2 µL of REST (Millipore, Cat. 17-641) Rabbit polyclonal antibody overnight at 4C Rabbit polyclonal antibody overnight at 4C. After de-crosslinking, ChIP DNA was purified using the Qiagen MinElute PCR Purification Kit (Qiagen, Cat. 28004); libraries for H3K4Me3, H3K27Me3, REST, and 1% Input were generated for NDC and *PSEN1*^*M146L*^ conditions at the UC San Diego IGM sequencing core using Illumina TruSeq LT adaptors (Illumina Cat. FC-121-2001). Sequencing was performed on an Illumina HiSeq4000 generating Single-Read, 75bp reads with an average of 20 million reads per sample.

ChIP-Seq data preprocessing was performed using the BBDuk package (v37.95) in the BBTools suite, removing sequencing adaptors and selecting for all -end reads above a quality score threshold (Phred Q>10). Trimmed ChIP-Seq reads were aligned to the GRCh38 human genome (GCA_000001405.15 with no alternative analysis) using BBMap v37.95 in the BBtools suite with the options maxindel=20 ambig=random, followed by sorting and indexing of bam files using Samtools v1.3, annotation of PCR duplicates using Picard v2.3.0 markDuplicates with the option VALIDATION_STRINGENCY=LENIENT, and removal of PCR duplicates using Samtools v1.3^57^ command view with the options -b -h -F 1024. ChIP-Seq peak calling and differential histone methylation was performed using Homer v4.10^28^; briefly, tags were generated from preprocessed bam files using the function makeTagDirectory, peaks called at a FDR of 0.05 using the function findPeaks with the options -style histone -gsize 2.7e9 -fdr 0.05 -region -size 500 for H3K4Me3 or -style histone -gsize 2.7e9 -fdr 0.05 -region -size 2000 for H3K27Me3. Differential histone methylation calculated using the function getDifferentialPeaks with the options -F 2 and -size 600 for H3K4Me3 and -F 2 and -size 2500 for H3K27Me3. REST peaks were called at a FDR of <0.05 using the function findPeaks with the options -style factor -gsize 2.7e9 -fdr 0.05 -region -size 250. The Homer function annotatePeaks was used to annotate REST bound genes or annotate regions of differential histone methylation to the GRCh38 genome as well as generate histograms of ChIP read density at histone methylation peak centers for genes with differential expression and differential histone methylation with the options -size 5000 -hist 50 for H3K4Me3 read density or -size 10000 -hist 100 for H3K27Me3 read density.

### Human Brain Transcriptomic Analysis

Microarray data corresponding to NDC and *PSEN1*^*M139T*^ EOFAD human brain samples from a previously published study^45^ were retrieved from the NCBI Gene Expression Omnibus (GEO) database (GSE39420) as raw CEL intensity files. 10 samples from this dataset, all obtained from the posterior cingulate cortex region, were selected for analysis: 6 non-demented control subject brains samples and 4 EOFAD brain samples from patients harboring the *PSEN1*^*M139T*^ mutation. Samples were background corrected and log_2_ normalized using the ‘RMA’ function^64–66^ in the *oligo* package v1.44.0^67^ in R v3.5.0. Differentially expressed genes were determined using the *limma* v3.36.2^68^ package, selecting for genes with a FDR-adjusted p-value (q-value) < 0.05.

### ISMARA Data Analysis

Quality and adaptor trimmed fastq.gz RNA-Seq files for NDC and *PSEN1*^*M146L*^ hiPSC-derived neurons as well as CEL microarray files for NDC and *PSEN1*^*M139T*^ EOFAD human brain samples were uploaded to ismara.unibas.ch^12^ for automated processing. For RNA-Seq and microarray experiments, sample averaging was performed following motif enrichment. In order to determine a directional z-score for each enriched motif identified, the z-score for each given motif was multiplied by the sign of the Pearson correlation for the TF associated with the motif and the direction of change in expression for the genes associated with each motif (−1 for downregulated genes, +1 for upregulated genes). For motifs associated with miRNAs, qPCR expression (miR-9, miR-124) or literature evidence was used to determine a positive of negative correlation with gene targets of the given motif.

### Transcription Factor Target, miRNA Target, and Histone Modification Enrichment

Transcription Factor (TF) target enrichment was carried out using the using the GSEAPreranked function in GSEA v3.0^8,69^ on all genes identified in each line and Enrichr^11,70^ to investigate the TFs potentially controlling genes found to be differentially expressed. For TF-target analysis we used the ENCODE/ChEA Consensus database, incorporating the consensus genes targets identified for 104 TFs from the Encyclopedia of DNA Elements (ENCODE) project ^71^ and Chip-X Enrichment Analysis (ChEA) datasets, and for miRNA-target analysis we used the miRTarBase v2017^72^ database. For Enrichr analysis, differentially up- or downregulated genes were enriched in an unweighted manner for each gene set library, generating an Enrichr Combined Score ranked list of TFs. For enrichments using the GSEAPreranked function, a pre-ranked list of genes was generated for each condition (*PSEN1*^*M146L*^ and *PSEN1*^*A246E*^ hiPSC-derived neurons, *PSEN1*^*M146L*^ human brain samples) by taking the sign of the log^2^ fold expression change relative to NDC multiplied by the –log_10_ of the p-value for all genes sequenced or probes in the microarray. This ranked list was used for all GSEAPreranked enrichments, using the options -rnd_seed timestamp -scoring_scheme classic -set_max 5000 -set_min 15. Custom GMT files were generated for the ENCODE/ChEA Consensus and miRTarBase v2017 databases by downloading each from Enrichr and converting into the GSEA GMT format. Further, we generated a custom TF-target list of endotype-associated TFs obtained from GEO (Table S1). Peaks for each TF were either downloaded from GEO provided by the original authors or raw sequencing data was uploaded to CRUNCH for peak identification. Peaks were annotated with the Homer v4.10 function annotatePeaks, and each list of target genes filtered for HGNC protein coding genes. For the MYT1L target gene list, a custom Homer motif file was generated from the MYT1L_200-623_Primary motif provided in Mall *et al.* 2017, motif locations identified using the Homer v4.10 function scanMotifGenomeWide, and subsequently annotated with annotatePeaks. This list of genes was first filtered for HGNC protein coding genes and then further filtered for differential gene expression upon MYT1L knockdown or overexpression presented in either Mall *et al.* 2017 or Yokoyama *et al.* 2014.

### Gene Set Enrichment and Pathway Analysis

Gene Set Enrichment and Pathway Analysis for RNA-Seq was performed primarily by two complementary approaches: 1) Gene Ontology (GO)^73^ and Reactome Pathway^74^ enrichment of differentially expressed genes using PANTHER v13.1^7^ and 2) pre-ranked Gene Set Enrichment Analysis using the GSEAPreranked function in GSEA v3.0^8,69^. For all enrichments using PANTHER, a list of differentially expressed genes with the corresponding log_2_ fold RNA expression was created for each comparison in question and a statistical enrichment test performed using the default PANTHER settings. Enrichments were performed using the expanded GO v1.2 Biological Process gene sets and Reactome Pathway v58 gene sets; enriched gene sets and pathways were ranked by the –log_10_ of the FDR-adjusted p-value. For all enrichments using the GSEAPreranked function, a pre-ranked list of genes was generated for each condition (*PSEN1*^*M146L*^ and *PSEN1*^*A246E*^ hiPSC-derived neurons, *PSEN1*^*M139T*^ human brain samples) by taking the sign of the log^2^ fold expression change relative to NDC multiplied by the – log_10_ of the p-value for all genes sequenced or probes in the microarray. This ranked list was used for all GSEAPreranked enrichments, using the options -rnd_seed timestamp -scoring_scheme classic -set_max 5000 -set_min 15. GMT files were downloaded from the MSigDB collections for all Hallmark v6.2, Gene Ontology, KEGG^75^, and Reactome Pathway gene sets. These GMT files were used for all GSEA Preranked gene set and TF enrichments. For gene set overrepresentation analysis on ATAC-Seq, ChIP-Seq, and *PSEN1*^*M139T*^ human brain samples, Metascape analysis was performed using GO – Biological Process, Reactome, and KEGG pathway datasets on the top 3000 significant DEGs (log_2_ fold-change > |0.5|, q-value < 0.01), differentially accessible chromatin (log_2_ fold-change > |0.5|, adj. p-value < 0.05), or differential histone methylation (log fold-change > |2|, adj. p-value < 0.05).

### miRNA qPCR assay

Total RNA from magnetically purified NDC, *PSEN1*^*M146L*^, and *PSEN1*^*A246E*^ (replicates, *n* = 3) human iPSC-derived neurons was prepared using miRNeasy Micro Kit (Qiagen Cat. 217084), based on manufacturer’s procedures. cDNA was produced using TaqMan Advanced miRNA cDNA Synthesis Kit (Life Technologies, Cat. A28007) according to the manufacturer’s instruction. Briefly, miRNA was polyadenylated at 37C for 45min using 20ng total RNA in a 5µL reaction. After heat-stop of the polyadenylation reaction at 60C for 10min, the reaction was held on ice for 2min. The adaptor was then ligated to the miRNAs at 16C for 1hr using RNA ligase in a 15µl reaction. Next, cDNA was synthesized from the adapter-ligated miRNAs. qRT-PCR for miRNAs were carried out using advanced TaqMan Advanced miRNA assay (Life Technologies, Cat. A25576), with TaqMan Fast Advanced Master Mix (Life Technologies, Cat. 4444557) and TaqMan prime probe sets for miR-124 (Hsa-miR-124-3P, 477879_mir) and miR-9 (Hsa-miR-9-5P, 478214_mir). The qRT-PCR program was as follows: 95C for 10min, leading to 40 cycles at 95C for 10min, followed by 60C for 20sec. The log_2_ expression fold change for each gene was calculated by the ΔΔCt method.

### Cell Cycle Assay

Magnetically purified NDC, *PSEN1*^*M146L*^, and *PSEN1*^*A246E*^ (replicates, *n* = 3) human iPSC-derived neurons were plated in 96 well plates. Cells were fixed in 70% ethanol and stored at 4C until the day of assay. Cells were stained with FxCycle™ Far Red Stain (Thermo Fisher, Cat. F10348) according to the manufacturer’s protocol. Stained cells were analyzed by flow cytometry using an Accuri C6 flow cytometer (BD Biosciences).

### Statistical analysis

GraphPad Prism v7.0e software was used for statistical analysis to calculate either student’s t-test or 1-way ANOVA with post-test analysis (Dunnett’s multiple comparison test) for Cell Cycle and qPCR assays.

### qPCR validation

Total RNA from magnetically purified NDC, *PSEN1*^*M146L*^, *PSEN1*^*A246E*^, and *PSEN1*^*H163R*^ (replicates, *n* = 3) human iPSC-derived neurons was prepared using RNeasy Plus Micro Kit (Qiagen, Cat. 74034) according to the manufacturer’s protocol. cDNA was synthesized from total RNA using random primer with iScript cDNA synthesis kit (BioRad, Cat. 1708890) according to the manufacturer’s protocol. qRT-PCR reactions were conducted using a CFX96 thermocycler (BioRad) with cDNA template, using TaqMan Fast Advanced Master Mix (Life Technologies, Cat. 4444557) and TaqMan primer probe sets (ThermoFisher). TaqMan assay ID numbers used: *GAD1* (Hs01065893_m1), *HDAC1* (Hs02621185_s1), *E2F1* (Hs00153451_m1), *CDK1* (Hs00938777_m1), *hes1* (Hs00172878_m1), *tp53* (Hs01034249_m1), *ccl2* (Hs00234140_m1), *CDKN1A* (Hs00355782_m1), and *PGK1* (Hs00943178_g1). The qRT-PCR program was as follows: 95C for 10min, leading to 40 cycles of 95C for 20sec, followed by 60C for 20sec. The log_2_ expression fold change for each gene was calculated by the ΔΔCt method.

### IPA Analysis

Ingenuity Pathway Analysis^46^ was performed on *PSEN1*^*M146L*^ and *PSEN1*^*A246E*^ hiPSC-derived neuron RNA-Seq DEGs and *PSEN1*^*M139T*^ human brain sample microarray DEGs, identifying positively and negatively z-score enriched causal upstream regulators (TFs).

---

**Fig. S1.**
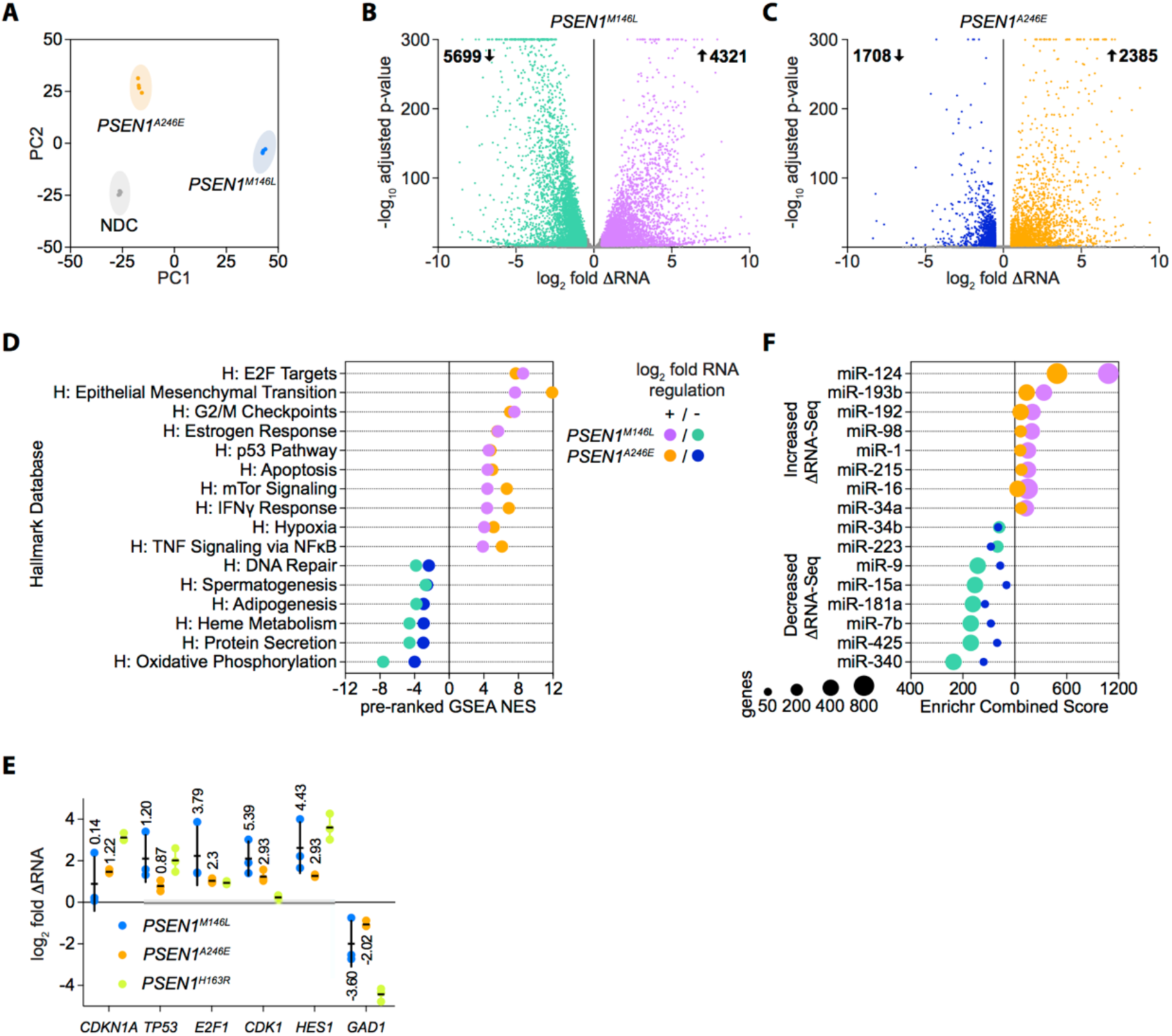
**A** Principal Component Analysis of NDC, *PSEN1*^*M146L*^, and *PSEN1*^*A246E*^ hiPSC-derived neurons (n = 4). **B-C** log_2_ fold change of differentially expressed genes in **B** *PSEN1*^*M146L*^ or **C** *PSEN1*^*A246E*^ hiPSC-derived neurons relative to non-demented control (NDC) with a False Discovery Rate (FDR) adjusted p-value (q-value) < 0.01 (n = 4). **D** Common pre-ranked GSEA normalized enrichment scores in *PSEN1*^*M146L*^ and *PSEN1*^*A246E*^ hiPSC-derived neurons using the Hallmark database (FWER < 0.01). **E** Confirmation of pathway-relevant differentially expressed genes by qPCR in *PSEN1*^*M146L*^, *PSEN1*^*A246E*^, or *PSEN1*^*H163R*^ hiPSC-derived neurons. (n=3) **F** Top enriched miRNA in the upregulated and downregulated *PSEN1*^*M146L*^ and *PSEN1*^*A246E*^ conditions by Enrichr using the miRTarBase v7.0 miRNA database (FDR adjusted p-value < 0.05).

**Fig. S2.**
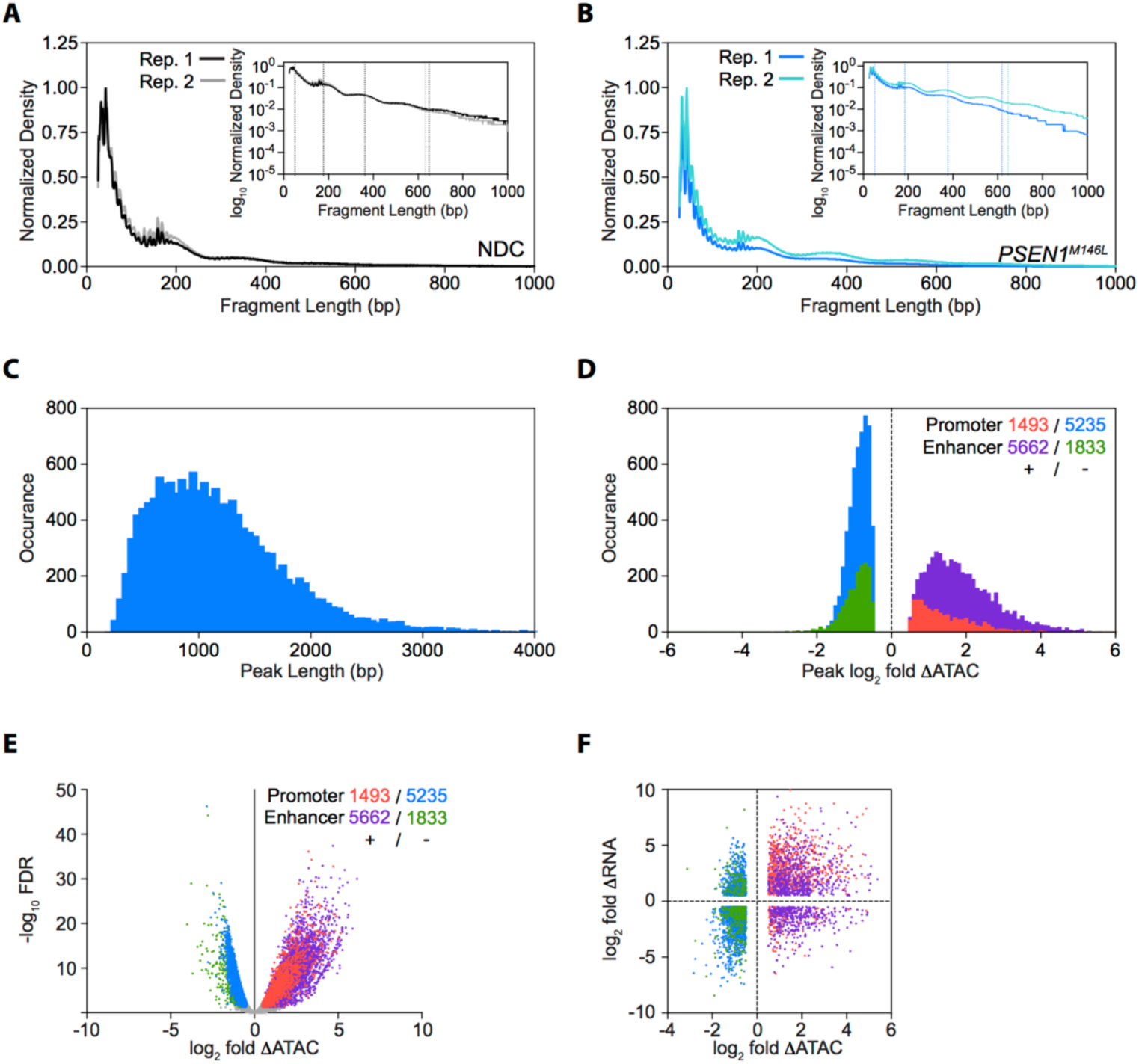
**A-B** Insert size distribution of ATAC-Seq insert sizes in **A** Non-Demented Control (NDC) and **B** *PSEN1*^*M146L*^ hiPSC-derived neurons. **C** ATAC-Seq differential peak width histogram in *PSEN1*^*M146L*^ hiPSC-derived neurons relative to NDC. **D-E** ATAC-Seq **D** differential peak log_2_ fold intensity histogram and **E** volcano plot diagram in *PSEN1*^*M146L*^ hiPSC-derived neurons relative to NDC, broken down by promoter-occurring and candidate enhancer regions.

**Fig. S3.**
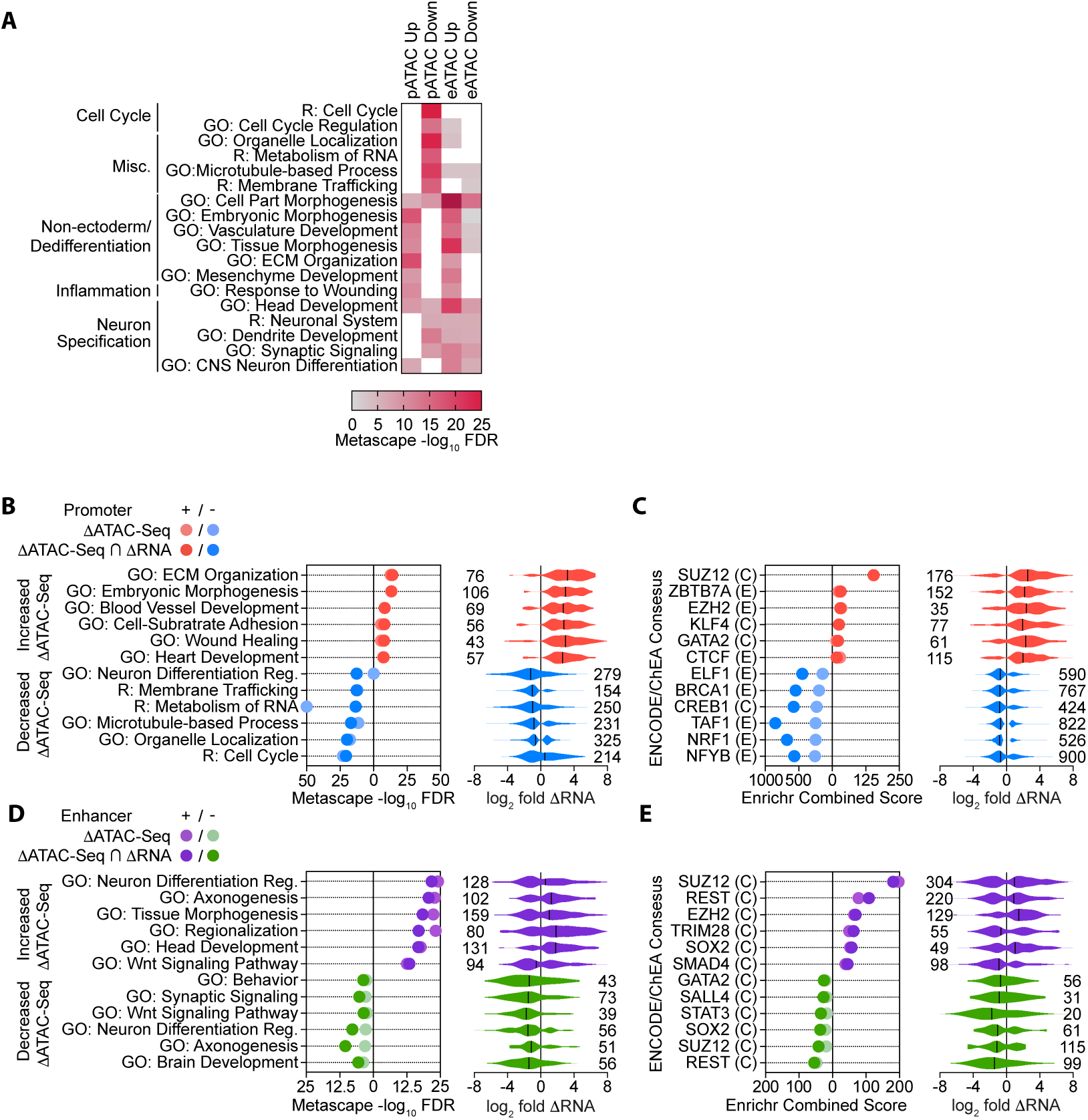
**A** Metascape enrichment heatmap of key endotype-related GO – Biological Process and Reactome gene terms in upregulated or downregulated regions of differentially accessible chromatin in the *PSEN1*^*M146L*^ condition occurring within the promoter region (pATAC) or outside the promoter region (eATAC). **B-C** Left, **B** Metascape and **C** Enrichr TF-target enrichment all differentially accessible promoter regions (light red, light blue) and differentially accessible promoter regions with differential gene expression (dark red, dark blue) using GO – Biological Process and Reactome (**B**) or Encode/ChEA Consensus (**C**) databases; right, violin plot of log_2_ fold change for significantly enriched gene terms for differentially accessible promoter regions with differential gene expression. **D-E** Left, **B** Metascape and **C** Enrichr TF-target enrichment all differentially accessible candidate enhancer regions (light red, light blue) and differentially accessible promoter regions with differential gene expression (dark red, dark blue) using GO – Biological Process and Reactome (**B**) or Encode/ChEA Consensus (**C**) databases; right, violin plot of log_2_ fold change for significantly enriched gene terms for differentially accessible promoter regions with differential gene expression.

**Fig. S4.**
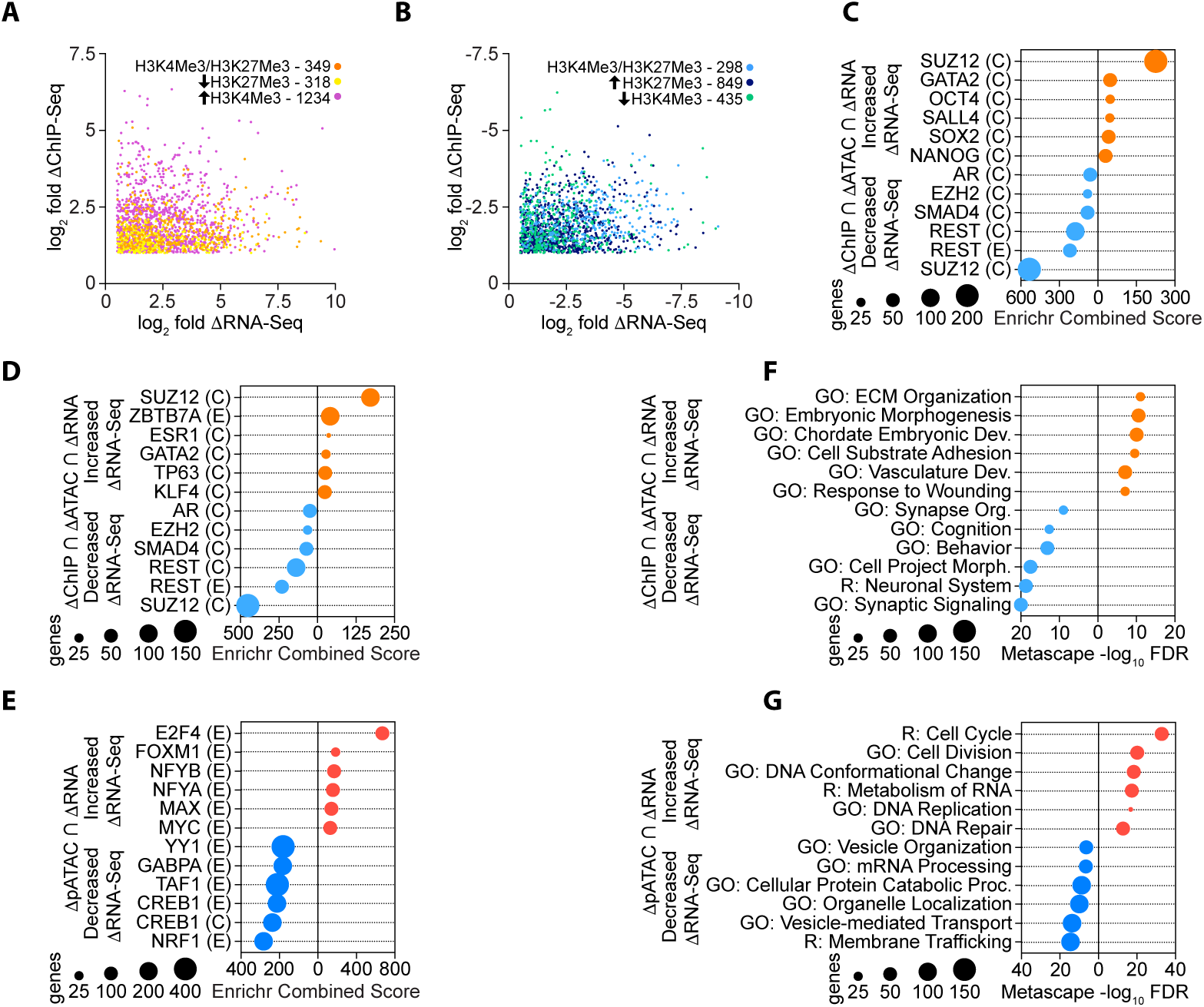
**A-B** log_2_ fold change in H3K trimethylation and RNA expression in genes with **A** increased gene expression and corresponding change in H3K trimethylation state (increased H3K4Me3 and/or decreased H3K27Me3) or **B** decreased gene expression and corresponding change in H3K trimethylation state (decreased H3K4Me3 and/or increased H3K27Me3). **C** Enrichr TF enrichment of genes with differential H3K trimethylation, chromatin accessibility, and gene expression using the ENCODE/ChEA Consensus database. **D-E** Enrichr TF enrichment of genes with **D** differential H3K trimethylation, chromatin accessibility, and gene expression or **E** differential chromatin accessibility in the promoter and gene expression alone using the ENCODE/ChEA Consensus database. **F-G** Metascape enrichment of genes with **F** differential H3K trimethylation, chromatin accessibility, and gene expression or **G** differential chromatin accessibility in the promoter and gene expression alone.

**Fig. S5.**
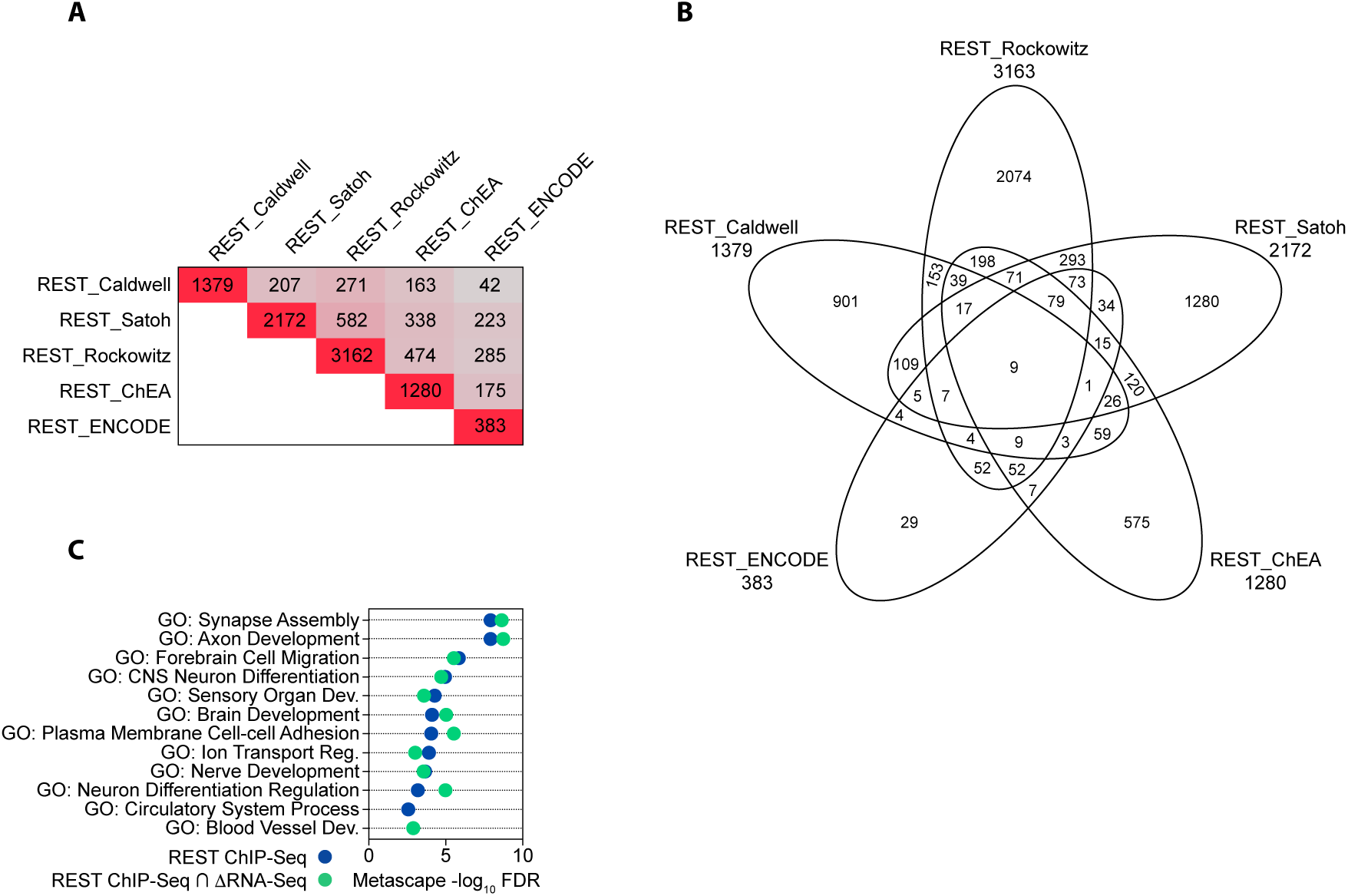
**A-B** Intersection of target genes identified by REST ChIP-Seq in *PSEN1*^*M146L*^ hiPSC-derived neurons compared with four other REST target gene sources: Enrichr ENCODE Consensus, Enrichr ChEA Consensus, Satoh *et al.* 2013, and Rockowitz *et al.* 2015. **C** Top Metascape enrichment of all REST ChIP-Seq bound genes and REST ChIP-Seq bound genes with differential gene expression in the *PSEN1*^*M146L*^ condition.

**Fig. S6.**
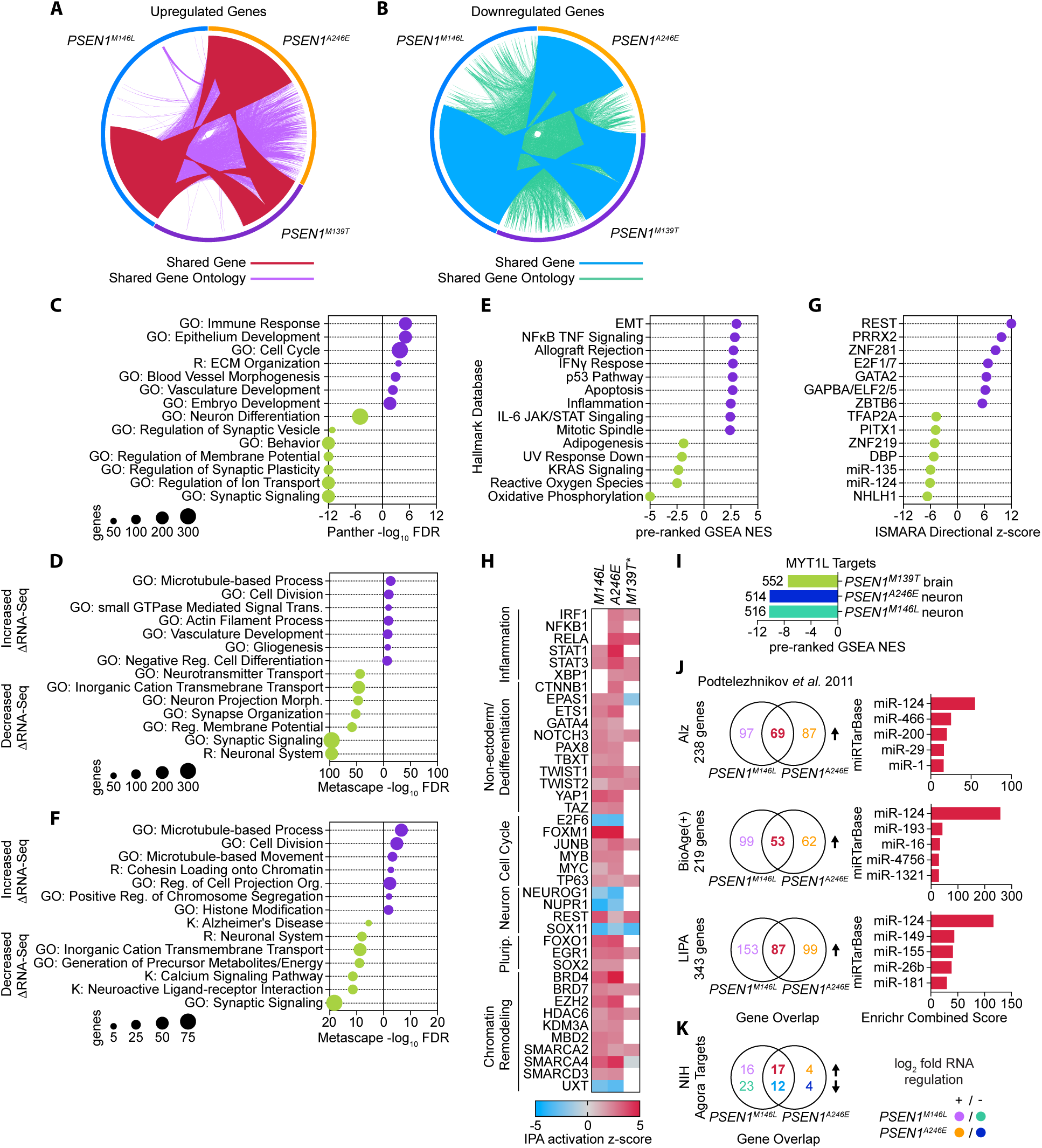
**A** PCA of NDC and *PSEN1*^*M139T*^ human brain sample microarray expression from Antonell *et al.* 2013; outlier NDC sample in red. **B-C** Circos plot of overlapping gene and genes with shared gene ontology between *PSEN1*^*M139T*^ human brain samples and *PSEN1*^*M146L*^ and *PSEN1*^*A246E*^ hiPSC-derived neurons for **B** upregulated genes and **C** downregulated genes. **D-E** Enrichment analysis of differentially expressed genes in *PSEN1*^*M139T*^ human brain samples using the GO – Biological Process and Reactome databases by **D** Panther or **E** Metascape enrichment. **F** Pre-ranked GSEA normalized enrichment scores in *PSEN1*^*M139T*^ human brain samples using the Hallmark database (FWER < 0.01). **G** ISMARA motif analysis in *PSEN1*^*M139T*^ human brain samples identifying the TFs and miRNAs with most significant motif-associated activity change based on ISMARA z-score and Pearson correlation. **H** Metascape enrichment analysis of unique DEGs in *PSEN1*^*M139T*^ human brain samples not differentially expressed in *PSEN1*^*M146L*^ and *PSEN1*^*A246E*^ hiPSC-derived neurons using the GO – Biological Process and Reactome databases. **I** Ingenuity Pathway Analysis activation z-score for transcriptional regulators in *PSEN1*^*M139T*^ human brain samples and *PSEN1*^*M146L*^ and *PSEN1*^*A246E*^ hiPSC-derived neurons, curated into key modulated disease-associated endotypes. **J** Pre-ranked GSEA normalized enrichment score for MYT1L target genes in *PSEN1*^*M139T*^ human brain samples and *PSEN1*^*M146L*^ and *PSEN1*^*A246E*^ hiPSC-derived neurons. K Overlap with genes upregulated in *PSEN1*^*M146L*^ or *PSEN1*^*A246E*^ hiPSC-derived neurons (left) and Enrichr miRNA enrichment (right) for Alz, BioAge(+), and LIPA sporadic Alzheimer’s Disease-associated metagene networks identified in Podtelezhnikov *et al* 2011. **L** Overlap between the 95 NIH Agora target genes identified from sporadic AD cases and upregulated or downregulated genes in *PSEN1* hiPSC-derived neurons.

